# Oscillatory entrainment mechanisms and anticipatory predictive processes in Autism Spectrum Disorder (ASD)

**DOI:** 10.1101/2020.05.07.083154

**Authors:** Shlomit Beker, John J. Foxe, Sophie Molholm

## Abstract

Anticipating near-future events is fundamental to adaptive behavior, whereby neural processing of predictable stimuli is significantly facilitated relative to non-predictable inputs. Neural oscillations appear to be a key anticipatory mechanism by which processing of upcoming stimuli is modified, and they often entrain to rhythmic environmental sequences. Clinical and anecdotal observations have led to the hypothesis that people with Autism Spectrum Disorder (ASD) may have deficits in generating predictions in daily life, and as such, a candidate neural mechanism may be failure to adequately entrain neural activity to repetitive environmental patterns. Here, we tested this hypothesis by interrogating rhythmic entrainment both behaviorally and electrophysiologically. We recorded high-density electroencephalography in children with ASD (n=31) and Typically Developing (TD) age- and IQ-matched controls (n=20), while they reacted to an auditory target as quickly as possible. This auditory event was either preceded by predictive rhythmic visual cues, or not. Results showed that while both groups presented highly comparable evoked responses to the visual stimuli, children with ASD showed reduced neural entrainment to the rhythmic visual cues, and altered anticipation of the occurrence of these stimuli. Further, in both groups, neuro-oscillatory phase coherence correlated with behavior. These results describe neural processes that may underlie impaired event anticipation in children with ASD, and support the notion that their perception of events is driven more by instantaneous sensory inputs and less by their temporal predictability.

## INTRODUCTION

The sensory environment provides us with context that both constrains and predicts near-future events, allowing for anticipatory planning and decision-making (Rao & Ballard, 1999; Lakatos *et al.*, 2008). The ability to anticipate relevant events is highly advantageous for adaptive behavior (Nobre *et al.*, 2007), leading to facilitated sensory processing of predictable inputs (Foxe & Simpson, 2005; Foxe *et al.*, 2005; Cui *et al.*, 2009; Kelly *et al.*, 2009; Cravo *et al.*, 2013). A recently forwarded hypothesis in autism research is that individuals with autism spectrum disorder (ASD) are impaired in their ability to use contextual information to modulate current processing (Mitchell & Ropar, 2004; Pellicano & Burr, 2012a), but see: (Brock, 2012)). This impairment stems from their altered use of prior experience to generate expectations about the environment (Pellicano & Burr, 2012b; Sinha *et al.*, 2014; Uljarevic *et al.*, 2017), and is associated with compromised adaptation processes (Lawson *et al.*, 2015). Indeed, in studies that involve perceptual judgments, individuals with ASD often do not calibrate neural sensitivity on the basis of the recent past, as is typically seen in controls (Turi *et al.*, 2015; Karaminis *et al.*, 2016; Turi *et al.*, 2016; Lieder *et al.*, 2019). While a common argument is that perceptual decisions follow Bayesian rules, in the sense that they are based on the prior likelihood of the events (Alais & Burr, 2004; Kersten *et al.*, 2004), people with ASD have altered perceptual judgment, where processing of priors may be underweighted. This underweighting of priors has been linked to impaired updating of events and impaired predictive processing in ASD (Pellicano & Burr, 2012a), and could explain prominent symptoms of the clinical phenotype of autism, such as reduced behavioral flexibility, insistence on sameness and rigidity of routines (Pellicano *et al.*, 2007; Pellicano, 2013; Van de Cruys *et al.*, 2014). Despite evidence for an imbalance in the influence of prior information versus sensory input on perception, the underlying neuronal processes remain poorly understood.

A wealth of research makes clear that neural entrainment, the phase-locking of brain oscillatory processes to rhythmic sensory inputs, is one prominent mechanism by which the brain encodes contextual cues to predict near-future events (Busch *et al.*, 2009; Schroeder & Lakatos, 2009; Gomez-Ramirez *et al.*, 2011; Fiebelkorn *et al.*, 2013; Gray *et al.*, 2015; Mazaheri *et al.*, 2018; Wilson & Foxe, 2020). According to this account, during entrainment to a predictable stimulus, the phase of slow oscillations aligns such that the stimulus presentation falls within an ‘optimal’ phase. These ‘excitability phases’ are thought to act as a sensory selection mechanism when oscillations are in the lower frequency delta and theta ranges (Schroeder & Lakatos, 2009). Given mounting evidence for impaired updating of priors in the face of changing environments in ASD, we considered that instantaneous tracking of information might be impaired. We hypothesized that impaired entrainment in individuals with ASD would lead to altered tracking of temporal regularities in the environment, and in turn, this would negatively impact anticipation of, and response to, temporally predictable events. Using a child-friendly novel paradigm, we generated temporal expectation through rhythmic visual stimuli that were informative of the timing of an upcoming auditory target event. We focused our analyses on entrainment of neural oscillations to the rhythmic visual cues, and on participants’ anticipation of the target, as indexed by the contingent negative variation (CNV) response, a negative deflecting EEG component indicating preparation for an anticipated target (Walter *et al.*, 1964; Dias *et al.*, 2003; Breska & Deouell, 2017). We expected that the extent of impairment of these neural processes would be related to target detection performance, since detection threshold and ongoing EEG phase have been shown to fluctuate together (Busch *et al.*, 2009; Vanrullen *et al.*, 2011; Lawson *et al.*, 2017). In addition, we tested whether the evoked responses to the auditory and visual sensory events were altered for the ASD group. Altered entrainment and anticipation, in conjunction with comparable evoked sensory potentials between ASD and controls would indicate dissociation between early sensory-based processing and high-order functions of tracking and anticipation. Alternatively, if both processes are impaired, this would suggest a more general impairment in neuronal functioning in response to sensory stimuli.

## METHODS

### Participants

31 children diagnosed with ASD and 20 IQ- and age-matched typically developing (TD) controls participated in the study (see Table 1).

**Table 1:** Mean ± SD of Age, sex and IQ of the two groups.

|  | Age | Sex | IQ (Full scale) | IQ (Non Verbal) | ADOS severity | RBS-R |
| --- | --- | --- | --- | --- | --- | --- |
| ASD | 7.88 ± 2.5 | 36 (M)<br>7 (F) | 93.27 ± 16.6 | 97.8 ± 15.1 | 7.65 ± 1.7 | 36.2 ± 26.9 |
| TD | 7.77 ± 1.45 | 7 (M)<br>12 (F) | 102.86 ± 9.2 | 93.4 ± 12.9 | NA | NA |

Participants ranged in age from 6 to 9 years old. Data of ASD participants were collected in the context of a study testing the efficacy of different behavioral interventions. The data presented are from the pretreatment session. ASD participants were recruited without regard to sex, race or ethnicity; TD participants were recruited to match the ASD group on age (two-tailed t-test for unpaired samples with unequal variances and α=0.05; t=-0.21; p=0.83). IQ quotients for performance (PIQ), verbal (VIQ), and full-scale (FSIQ) intelligence were assessed in all of the ASD, and a subset of the TD (10 out of 20) participants using the Wechsler Abbreviated Scales of Intelligence (WASI; (Stano, 1999)). Comparison of non-verbal IQ shows no significant differences between the groups (t-test for unpaired samples with unequal variances and α=0.05; t=-0.8; df=40; p=0.38). To be considered for the ASD group, participants had to meet diagnostic criteria for ASD on the basis of the following measures: (1) Autism Diagnosis Observation Schedule (ADOS-2); (2) Diagnostic Criteria for Autistic Disorder from the DSM-5; (3) Clinical impression of a licensed clinician with extensive experience in diagnosis and evaluation of children with ASD. Repetitive Behavior Scale-Revised (RBS-R) (Lam & Aman, 2007) questionnaire was collected to obtain continuous measures of ASD characteristics related to insistence on sameness such as ritualistic/sameness behavior, stereotypic behavior, and restricted interests. In order to be included in the TD group, participants had to have no history of neurological, developmental or psychiatric disorders or first degree relatives with a diagnosis of ASD, and had to be in age-appropriate grade at school.

Participants received nominal recompense for their participation (at $15 per hour for the TD participants and a total of $250 for participation in the treatment study for ASD participants). Exclusionary criteria for both groups included epilepsy, or premature birth (<35 weeks, with the exception of one ASD participant born at 32 weeks). Participants from both groups scored nonverbal IQ>80, with the exception of two ASD participants who scored 64 and 79 respectively. All participants passed a screen for normal or corrected-to-normal vision and normal hearing on the day of testing. Parents and/or guardians of all participants provided written informed consent. All procedures were approved by the Institutional Review Board of the Albert Einstein College of Medicine.

### Stimuli and Task

Visual stimuli were presented on a 25” ViewSonic screen (refresh rate: 60 Hz, pixel resolution: 1280×1024×32) of a Dell computer using Presentation^©^ software (Version 20.0, Neurobehavioral Systems, Inc., Berkeley, CA, www.neurobs.com). The stimuli consisted of a cartoon dog’s face image located at the center of the screen, and subtended ∼4.4° of visual angle. Auditory stimuli were delivered at an intensity of 75 dB SPL via a single, centrally-located loudspeaker (JBL Duet Speaker System, Harman Multimedia). The task was designed to test the hypothesis that children with ASD do not use temporally predictive information in a typical way, and that this is related to impaired entrainment. The task included two conditions: For the *Cue* condition, participants were presented with a sequence of 4 visual isochronous stimuli for a duration of 80ms each and an Stimulus Onset Asynchrony (SOA) of 650ms, followed by an auditory stimulus, presented 650ms after the last cue for a duration of 80ms. For the *No-Cue* control condition, the auditory stimulus was not preceded by the sequence of visual stimuli. Each target appeared 2600ms after the beginning of trial, during which participants were focused on the screen. In all other respects, the paradigm, including the timing of the stimuli, was identical between the conditions (See Fig. 1 for paradigm illustration). Cue and No-Cue conditions were blocked, and presented in runs of 25 trials per block, over a total of 20 blocks. The order of blocks within the experiment for a given participant was chosen randomly prior to each experimental session. Participants were seated at a fixed distance of 65 cm from the screen. In all trials, they were instructed to press a button on a response pad (Logitech Wingman Precision Gamepad) as soon as they heard the auditory tone. Responses occurring between 150-1500ms following the auditory tone were considered correct, and positive feedback was provided via presentation of a cartoon dog image and an uplifting sound. If the response was outside this time window, a running dog cartoon with a sad sound was presented to indicate that the response was too fast, and a sitting dog image with the sad sound was presented to indicate that the response was too slow (see Fig. 1). Frequent breaks were given as needed to ensure maximal task concentration.

**Figure 1:**
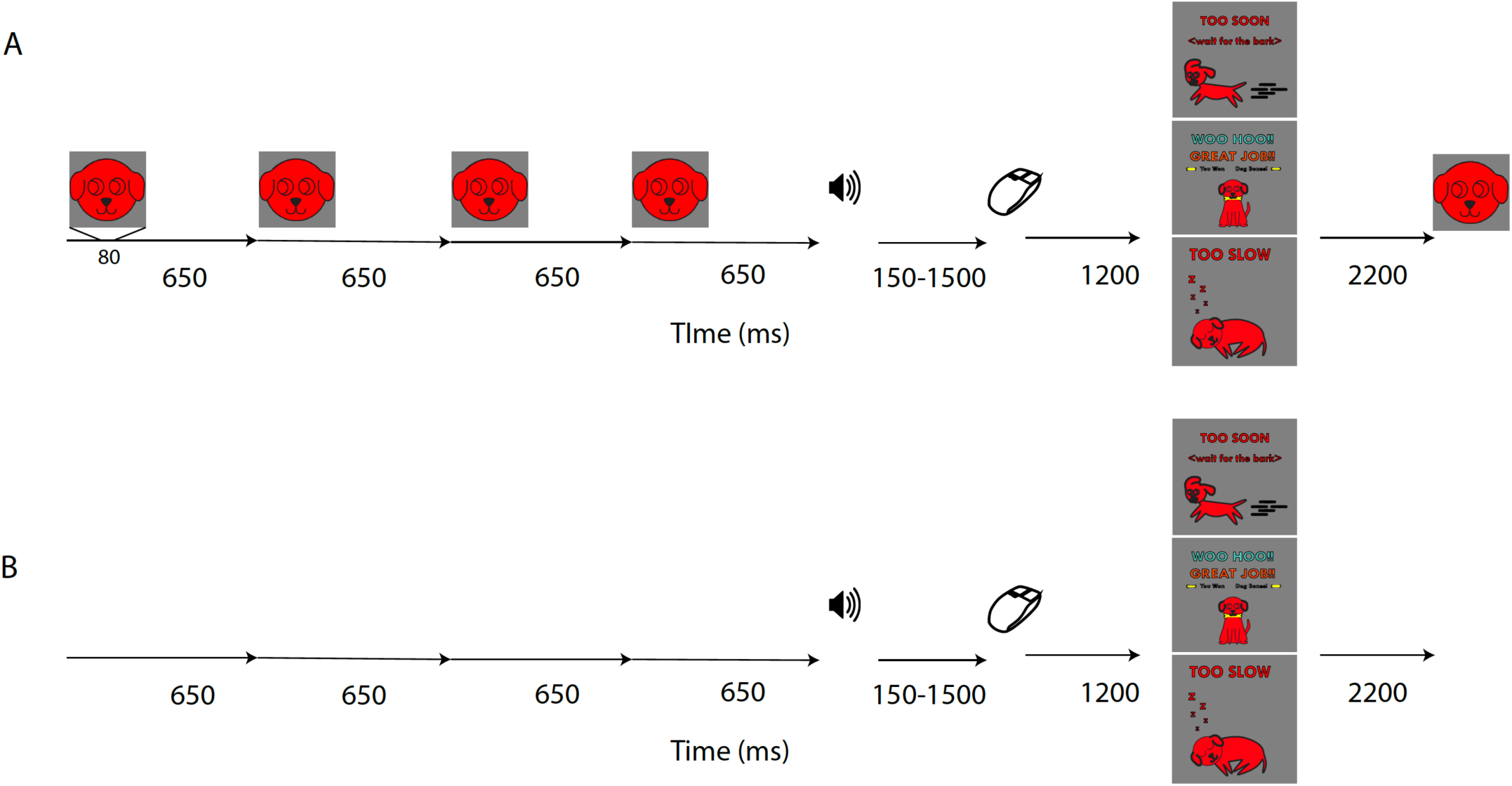
Schematic of Experimental Paradigm. **A)** Cue condition; **B)** No-Cue condition (identical in timing to Cue condition except that visual cues were not presented).

### Data acquisition

Response times were recorded with the Presentation^©^ software. EEG recordings were collected from 70 active channels (64 scalp channels and 6 external electrodes) at a digitization rate of 512 Hz, using ActiView (BioSemiTM, Amsterdam, The Netherlands). Analog triggers indicating the latencies of stimulus onsets and button presses were sent to the acquisition PC via Presentation® and stored digitally at a sampling rate of 512 Hz, in a separate channel of the EEG data file.

### Data processing

Data were processed and analyzed using custom MATLAB scripts (MATLAB r2017a, MathWorks, Natick, MA), and the FieldTrip toolbox (Oostenveld et al., 2011, Donders Institute for Brain, Cognition and Behavior, Radboud University, the Netherlands). A minimum number of 50 EEG trials per condition per analysis was set as a criterion for a participant to be included in the analysis, however, most participants had more than 100 trials in each condition (mean±sem TD: 168±37; ASD: 136±24). Due to occasionally long reaction times in some of the participants, only responses within 1000ms after stimulus presentation were considered as valid for further analysis of the behavior data.

### EEG Processing

To tailor analyses to each of our research questions, data processing involved four different sub-pipelines, as elaborated below. All epochs, except for those used for measuring entrainment (see 3 below) were down-sampled to 256 Hz, epoched and band-pass filtered between 0.1 and 55 Hz using Butterworth Infinite Impulse Response (IIR) windowing, with filter order of 5. Epochs were then de-meaned to normalize for DC shifts, and baseline-corrected, as specified below.

1. Visual Evoked Response (VEP): continuous data were epoched with respect to each visual stimulus: 200 ms before and 850 ms after stimulus onset, and baselined to the 100ms window before the onset of each visual stimulus. Two averaging approaches were taken, so that we could measure and compare between groups on a) the overall VEP, and b) sequence effects on the CNV, as follows:

a. VEP: To compare sensory-evoked potentials between the groups, trials were averaged across all stimuli, regardless of their placement in the sequence. Data were referenced to a frontal channel (AFz) to optimize visualization and measurement of the VEP over occipital scalp at channels O1, O2, and Oz. Comparison between the groups on the VEP was calculated on the voltage at the peak of the P1 component (∼100ms) for each participant.
b. CNV: To find differential patterns of anticipation with respect to the place of a cue in the sequence, trials were averaged separately for visual stimuli 1 through 4 in the trial. ERP analyses were focused on preparatory effects (the CNV) that are commonly seen over fronto-central sites (Walter *et al.*, 1964; Gaillard, 1976; Breska & Deouell, 2017), and were measured here at frontal channels (AFz, FP1, AF3, and AF4). To optimize visualization and measurement, data were referenced to the two ground electrodes: Common Sense Mode (CMS), and Driven Right Leg (DRL), used in BioSemi as active and passive electrodes respectively. CNV activity was measured prior to the upcoming stimulus in the sequence, at −200 to 0 ms relative to stimulus (visual or auditory) onset.
2. Entrainment: to measure entrainment of neural oscillations to the event timing, the EEG data were epoched at 3100ms before and 300ms after the auditory event, to encompass the full sequence of stimuli comprising a trial, and trials were baselined to the 100ms before the onset of the first visual stimulus in the sequence. Data were referenced to channel AFz. Analyses of frequency and phase were performed on epochs that were low-pass filtered at 55Hz, and high pass filtered at 0.1Hz. To visualize entrainment of oscillatory activity to the rhythm of visual events (1.5 Hz), data were low-passed filtered at 1.9 Hz. For this, a Finite Impulse Response (FIR) filter, with normalized passband frequency of 1.9 Hz, stopband frequency of 4 Hz, passband Ripple of 1Hz and stopband attenuation of 60 dB, was applied.
3. Auditory Evoked Response (AEP): To analyze evoked responses to the auditory target stimulus and possible differences between the groups with respect to condition, data were epoched 300ms Sec. before and 850ms Sec. after the auditory stimulus onset, and baselined to 100ms before auditory stimulus onset, and as per convention, referenced to a channel near the left mastoid (TP7). Statistical analysis was performed on data from fronto-central channels at the maximal peak of each participant’s P1 (∼50ms), N1 (∼100ms) and P2 (∼200ms) components.

For all pipelines, after epoching, a two-stage automatic artifact rejection was applied at the single trial level. First, channels that varied from the mean voltage across all channels and from the auto-covariance by 1 standard deviation were classified as bad. A maximum of six bad channels was set as an inclusion criterion for trials to be analyzed. For these trials, channels were interpolated using the nearest neighbor spline (Perrin *et al.*, 1987; Perrin *et al.*, 1989). Second, a criterion of ±120μV was applied. Electrodes that exceeded this criterion were considered bad. Participants that had less than 50 trials per condition were excluded from further analysis. Overall, 3 ASD and 1 TD participants were excluded from analysis due to insufficient number of trials.

### Analytic approach

Analysis of sensory event related potentials (ERPs) was conducted on visual (P1) and auditory (P1, N1, P2) components, at the relevant scalp sites (see above pipelines). For each participant, windows of 20, 40 and 60ms were centered at each of the components (at 50ms, 100ms, and 200ms respectively) as the area of interest. Then, the maximal (for P1, P2) or minimal (for N1) voltage was found for each participant within each of these windows.

For the anticipatory ERP analysis (i.e. the CNV), a window of interest was defined between −200 and 0 ms with relation to an expected stimulus. Voltages in this time window were averaged for every participant for each condition. To compare CNV amplitude differences between the groups in the Cue condition, we performed a two-tailed t-test with unpaired samples and unequal variances, and α=0.05. To analyze AEP amplitude differences between the groups, we performed 2-way ANOVA with α=0.05 and Group and Condition as factors. To compare CNV between the groups, we performed a 2-way ANOVA with α=0.05 and group and sequential place as factors.

In addition to the above predefined components for statistical analysis, an exploratory approach was taken to explore and visualize effects in other time windows and scalp regions. For this, we tested the entire data matrix. Statistical cluster plots (SCPs; (Molholm *et al.*, 2002) were generated by calculating paired two-tailed t-tests between the groups, in the VEP and the AEP, generated at each time point for each group, and across all channels. Time/channels points that meet the criterion of at least 10 consecutive time points and 2 consecutive channels with α=0.05, were marked as significant.

In the entrainment analysis pipeline (pipeline 2 above), Inter-trial Phase Coherence (ITPC) was calculated separately for each individual participant, between all trials in the two experimental conditions (cued and non-cued). This measure quantifies the consistency of the phase of responses across trials. ITPC was calculated as the circular variance of the phase across trials (Luo & Poeppel, 2007) at each frequency and time bin, and averaged across time bins. Coherence was calculated as following:

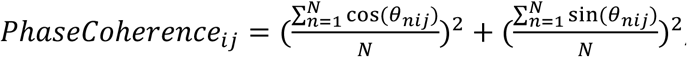, where *θ*_*nij*_ is the phase at frequency bin i and temporal bin j. Phase Coherence was calculated for the range of 0.6-2.7 Hz. A larger ITPC index corresponds to stronger coherence. The average number of trials per participant used to calculate ITPC for TD/ASD groups (with cue) was 195±9.9 and 158±7.6, respectively. ITPC was first calculated for the frequency domain, collapsed across the 3500ms. time window (including the four visual cues and auditory target) at all scalp electrodes. The frequency range 0.6–2.7 Hz was divided into 28 bins. To evaluate differences in ITPC between the groups, a 2-way ANOVA was performed on the ITPC values with Group and Cue as factors. In addition, to evaluate the time course of the ITPC, we calculated ITPC for each time and frequency bin in the range of the whole trial (2900ms), and at 2-7 Hz, respectively. For this, we used wavelets ranging from 2 to 7 Hz with 1 Hz width, and divided the time window of 400 to 2400ms post the first cue onset, in steps of 4ms. The edges of the time domain were trimmed due to requirements of wavelet time-frequency decomposition. The large frequency range was included in order to explore effects in higher frequencies than that of stimulation. For visualization, differences in ITPC between TD and ASD groups were calculated by subtracting ITPC values of ASD group averages from those of TD group averages, for each time and frequency point. Nonparametric statistics were computed using permutation tests. To assess the null distribution, the group labels were randomly intermixed for each subject, and the ITPC difference was computed with p<0.05 as a threshold. This procedure repeated 10000 times for ITPC across, on each frequency and time point in the 6*501 matrix (Zoefel *et al.*, 2017).

To test for correlations between behavior and measurements of entrainment, a linear Pearson correlation was performed between reaction times (RT) and ITPC. In addition, we considered standard deviation of RT (SD-RT), to test whether it might correlate better with variability of entrainment. SD-RT*ITCP correlation was calculated for each group individually, and across the two groups. To find if the distribution of the data points on the two axes mapped onto separate clusters that were dominated by one or the other participant group, K-means classification was performed on all participants SD-RT*ITPC correlation data points. K-means clustering partitions *n* samples into *k* clusters of greatest possible distinction. Each object belongs to the cluster with the nearest mean. The objective of K-Means clustering is to minimize total intra-cluster variance, or, the squared error function:

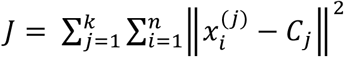, where *k* is the number of pre-defined clusters, *n* is number of samples, and *C* is the centroid of a cluster. Two centroids were defined for the combined dataset, using the elbow method (Ketchen & Shook, 1996). Evaluation of k-mean clustering was achieved using the Silhouette method, with squared Euclidean distance (Rousseeuw, 1987).

To assess potential associations between ASD severity and oscillatory entrainment, Pearson correlations between scores of behavioral/cognitive assessments and ITPC were tested for the ASD children. For behavioral/cognitive assessments, we used scores from Autism Diagnostic Observation Schedule (ADOS); Performance IQ, Verbal IQ and Repetitive Behavior Scale-Revised (RBS-R) questionnaire. P values were corrected for multiple comparisons, using Holm-Bonneferoni False Discovery Rate (FDR) method (Aickin & Gensler, 1996)

## RESULTS

We investigated the influence of an isochronous and predictable repeating stimulus on neural entrainment and anticipation processes in children with a diagnosis of ASD, using measurements of reaction times and of EEG dynamics.

### Behavioral Results

Reaction times (RT) were measured as the time between the onset of the auditory target and the participant’s motor response. Overall, both groups of participants had shorter latencies in the Cue compared with the No-Cue condition (Cue: 331±23 ms; No-Cue: 382±23 ms; F=5.8; d=1; p=0.018). In addition, there was a main effect of group, where the average RT was slower for ASD compared to TD participants, across conditions (ASD Cue: 396±24 ms; No-Cue: 465±24 ms; TD Cue: 331±23 ms; No-Cue: 382±23 ms; F=8.7; d=1; p=0.004). No-Cue*Group interaction was present (F=0.11; df=1; p=0.7; See Figure 2 (panels A and B are plotted with the toolbox available in (Ho *et al.*, 2019)), suggesting that both groups benefited equally from the presence of the cueing stimulus sequence.

**Figure 2:**
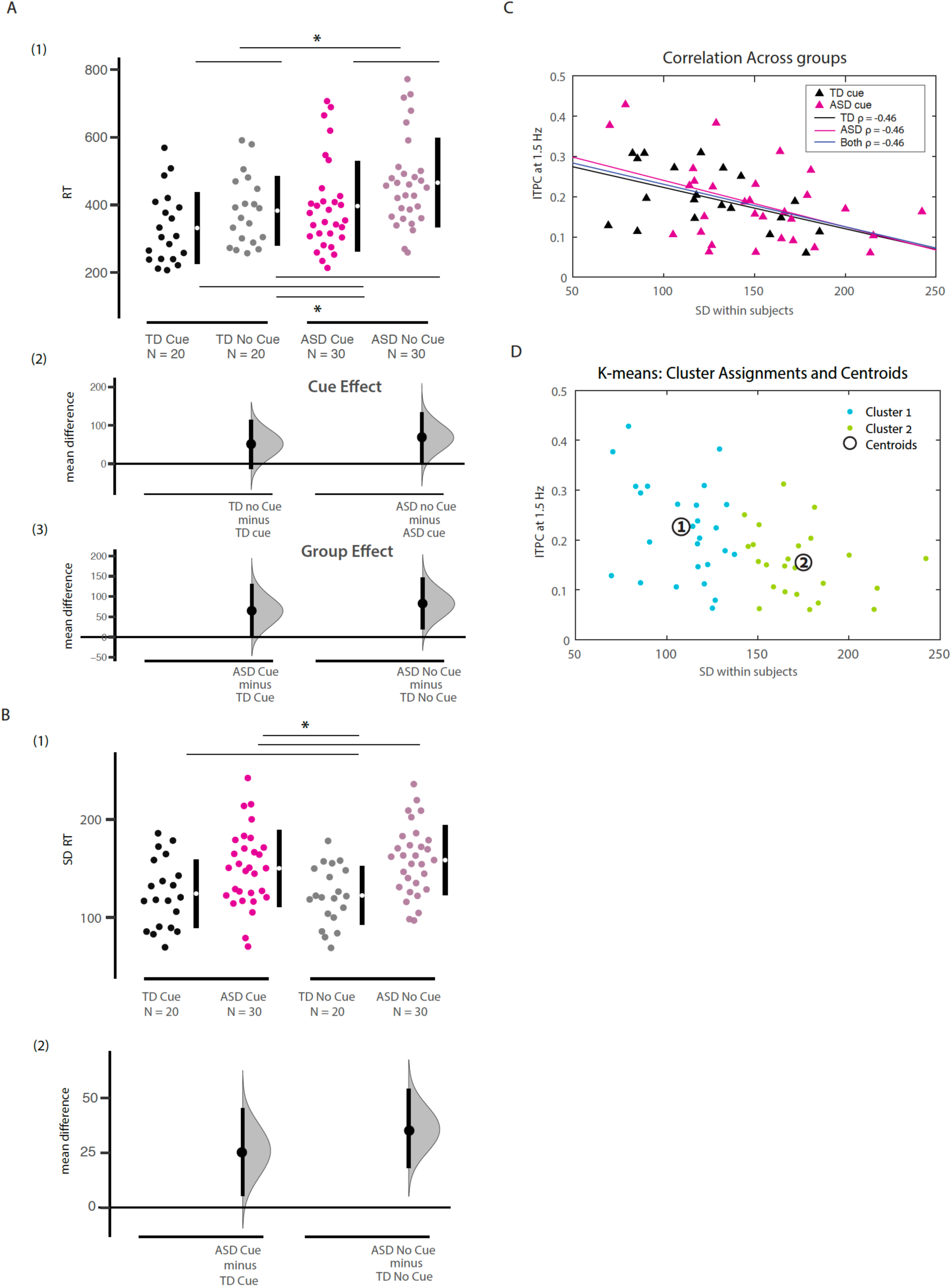
Behavioral results. **A)** Reaction times (RT) of participants in each group and condition. RTs of the ASD group (Cue: 396; No Cue: 465) were longer than those of TD group (Cue: 331; No Cue: 382), and faster for the Cue compared to the No Cue condition. There was no Group*Condition interaction. A1) Individual subject data; A2) RT difference between Cue and No Cue for each of the groups; A3) RT difference between groups for Cue (left) and No Cue (right) conditions. **B)** Standard Deviations (SD) of RTs. SD-RTs were higher for the ASD group compared to the TD group. B1) Individual subject data; B2) The difference between Cue and No-Cue conditions. **C)** The correlation between Inter-Trial Phase Coherence (ITPC) and SD-RT for each group separately, and when collapsed. **D)** K-means clustering on the basis of ITPC and SD-RT data with two centroids. Cluster 1 (Lower SD-RT/higher ITPC): 54% TD; Cluster 2 (Higher SD-RT/lower ITPC cluster): 75% ASD. 66% of ASD participants and 30% of TD participants fell into Cluster 2.

To measure differences in variability in behavior, the SD-RT was calculated for each individual participant and condition, and compared between the groups. On average, RTs of ASD participants were more variable than those of TD participants (SD-RT mean±SEM for the Cue condition: 120±14ms for TD and 178±18ms for ASD; and for the No-Cue condition: 122±13ms for TD and 187±17ms for ASD). There was a significant main effect of the factor Group (F=12.4, df=1; p<0.001) but not of Cue (F=0.2, df=1; p=0.7), and the Group*Condition interaction was not statistically significant (F=0.04; p=0.8).

### EEG Results

To account for the effect of introducing isochronous visual cues on different aspects of the brain responses, results are reported for the different pipelines used for the analysis (see above).

1. a) VEP: The visual evoked potential was measured at occipital sites (O1, Oz and O2) at 100ms after visual stimulus onset (P100). Mean VEPs did not differ significantly between the two groups (t-stat=-0.65; df=45, p=0.5). To test the robustness of seemingly different potentials at a later time window (400-600ms), where lower amplitudes for ASD than TD at frontal, central and parietal sites were observed (See Fig. 3A), we performed an exploratory two-tailed t-test for unpaired data (For occipital channels: t-stat=-2.02; df=45; p=0.04).

**Figure 3:**
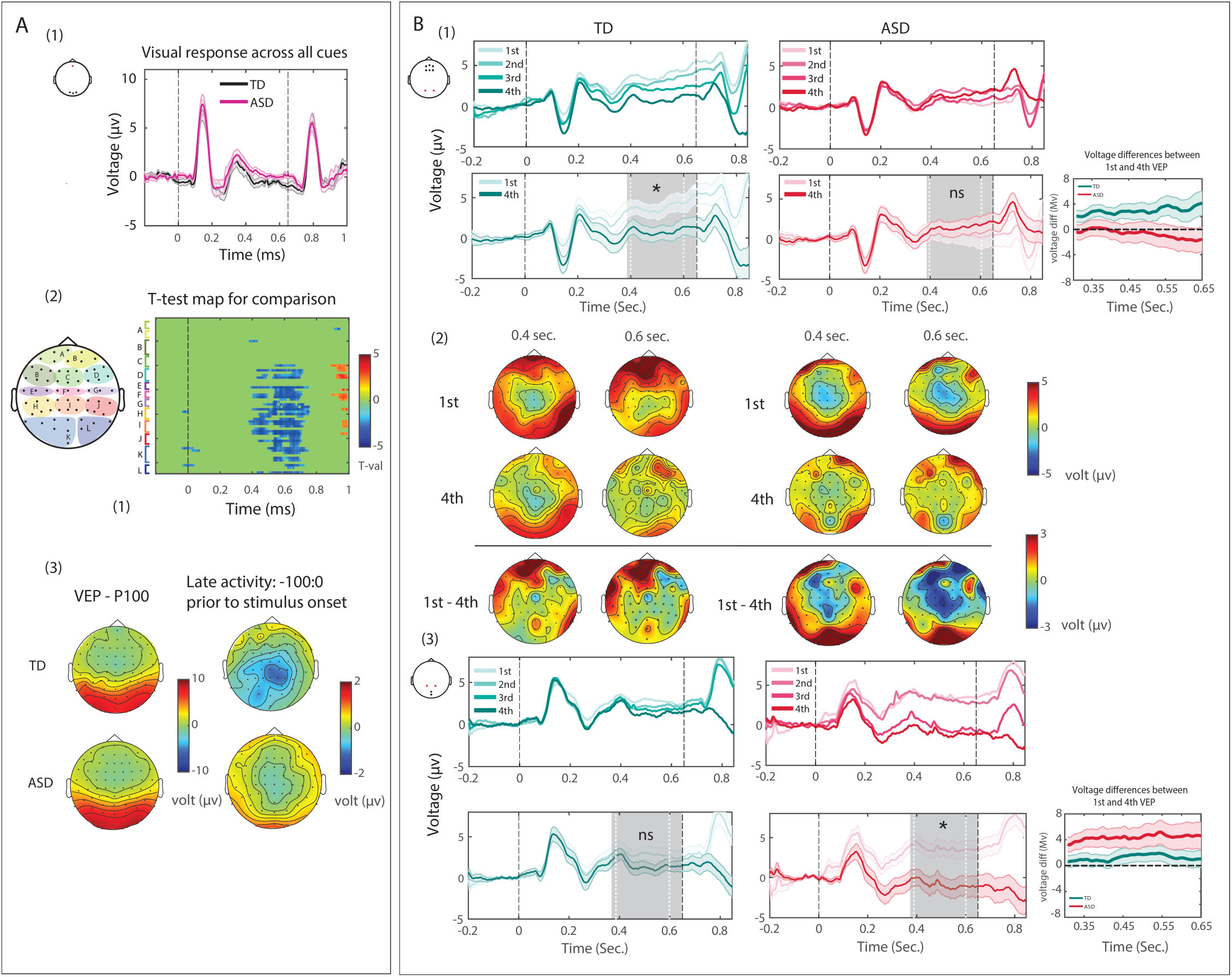
**A)** Visual evoked potential (VEP), averaged across all 4 cues. A1) Waveforms from occipital scalp for TD and ASD groups. A2) Statistical cluster plots of t-test results for ASD versus TD comparison for all channels and time points (plotting results where at least 10 consecutive time points were significant, for two adjacent channels). A3) Scalp voltage topographic maps for the P1 and the CNV. **B)** Anticipatory activity over frontal scalp channels: B1) Top-For each of the 4 sequential visual cues; Bottom-For the 1st and last (4^th^) visual cue. A graded anticipatory response, highlighted by gray, is present for the TD, but not the ASD, group. B2) Scalp topographic maps for the 1st and 4^th^ stimulus and the difference between them, at two time points prior to the upcoming stimulus. B3) Anticipatory activity over occipital scalp. Top: for each of the 4 sequential visual cues; Bottom: for the 1st and last (4^th^) visual cue. As highlighted by gray, a difference is apparent for the ASD, but not the TD, group. b) CNV: Considering the ERP to each stimulus in the sequence separately, a gradation of the CNV response is seen for the TD group, where voltage is more negative going the more advanced (later) the cue is in the sequence, in the time window of −200 to 0 ms with respect to stimulus onset. This gradation is best seen at frontal sites in TDs, where we focused our analysis (Fig. 3B). This negative-going change in voltage is not evident in the ASD group. For the ANOVA, we used Group and Cue order as factors, and included data from ERPs prior to the 1^st^ and the 4^th^ stimulus in each of the groups. We found a significant Group*Cue order interaction (F=4.8; df=1; p=0.03). No main effect for Group (F=2.5; df=1; p=-.12) or Cue order (F=0.2; df=1; p=0.64) was found. Post-hoc test on the interaction effect revealed a significant difference between voltage traces prior to 1^st^ and 4^th^ cues in the sequence of the TD group (p=0.04) but not the ASD group (p=0.97). Topographic plots show differential patterns for anticipatory activity between the groups. While the TD group shows clear anticipatory activity during the 200ms window prior to the stimulus over fronto-central scalp, the ASD group shows enhanced anticipatory activity over occipital regions (Fig. 3B). To test the apparent group difference in the topography of the graded anticipatory effects we first performed a 3-way ANOVA with Group, Cue order and Scalp region, including both frontal and occipital regions (Oz and Iz), as factors. This revealed a significant main effect for Cue order (F=5.5; df=1; p=0.02), and a Cue order*Group*Scalp region interaction (F=5.54; df=1; p=0.019) indicating that each group showed a different scalp focus for the graded anticipatory effects.
2. Entrainment: To visualize entrainment to the rhythm of stimulation (1.5 Hz), each epoch was individually low-pass filtered at 1.9Hz (see Methods). Then, epochs were averaged across participants. Broadband and low-pass filtered signals are shown in Fig. 4A. Fast Fourier Transform (FFT) analysis did not show between group differences in the maximal power at the stimulating frequency, 1.5Hz (Wilcoxon rank test zVal = −1.14; ranksum=460, p=0.25). To measure neural entrainment to the rhythm of cue presentation (1.5 Hz), the average phase across cue onsets was calculated for each individual trial at the stimulation frequency of 1.5 Hz. Polar plots show higher phase concentration for the TD group (Rayleigh test for non-uniformity p<0.001), compared with the ASD group (Rayleigh test for non-uniformity P=0.1; Fig. 4B). To measure coherence of phases across trials, Inter-Trial Phase Coherence (ITPC) was calculated across epochs for each participant and channel, and across participants in each group, for frequencies < 2.5Hz. ITPC is stronger over occipital (Oz, Iz) and parieto-occipital channels (PO3, POz, PO4) for the TD compare with ASD group (Fig. 4C). To evaluate differences in ITPC between the groups, a 2-way ANOVA was performed on the ITPC values with Group and Condition as factors. Main effects were found for Group (F=9.3; df=1; p=0.003) and Condition (F=8.6; df=1; p=0.004). There was no Group*Condition interaction (F=0; df=1; p=0.98). When calculated on each time and frequency points in the arrays, a strong difference in ITPC is seen between the groups in vicinity of cue onset times (Fig. 4E). Permutation tests (N=10000) were performed for every time and frequency point between ITPC of ASD and TD. As a correction for multiple comparisons, time/frequency points that meet the criterion of 10 consecutive points with α=0.05 were marked as significant (Fig. 4E) (See (Guthrie & Buchwald, 1991)).

**Figure 4:**
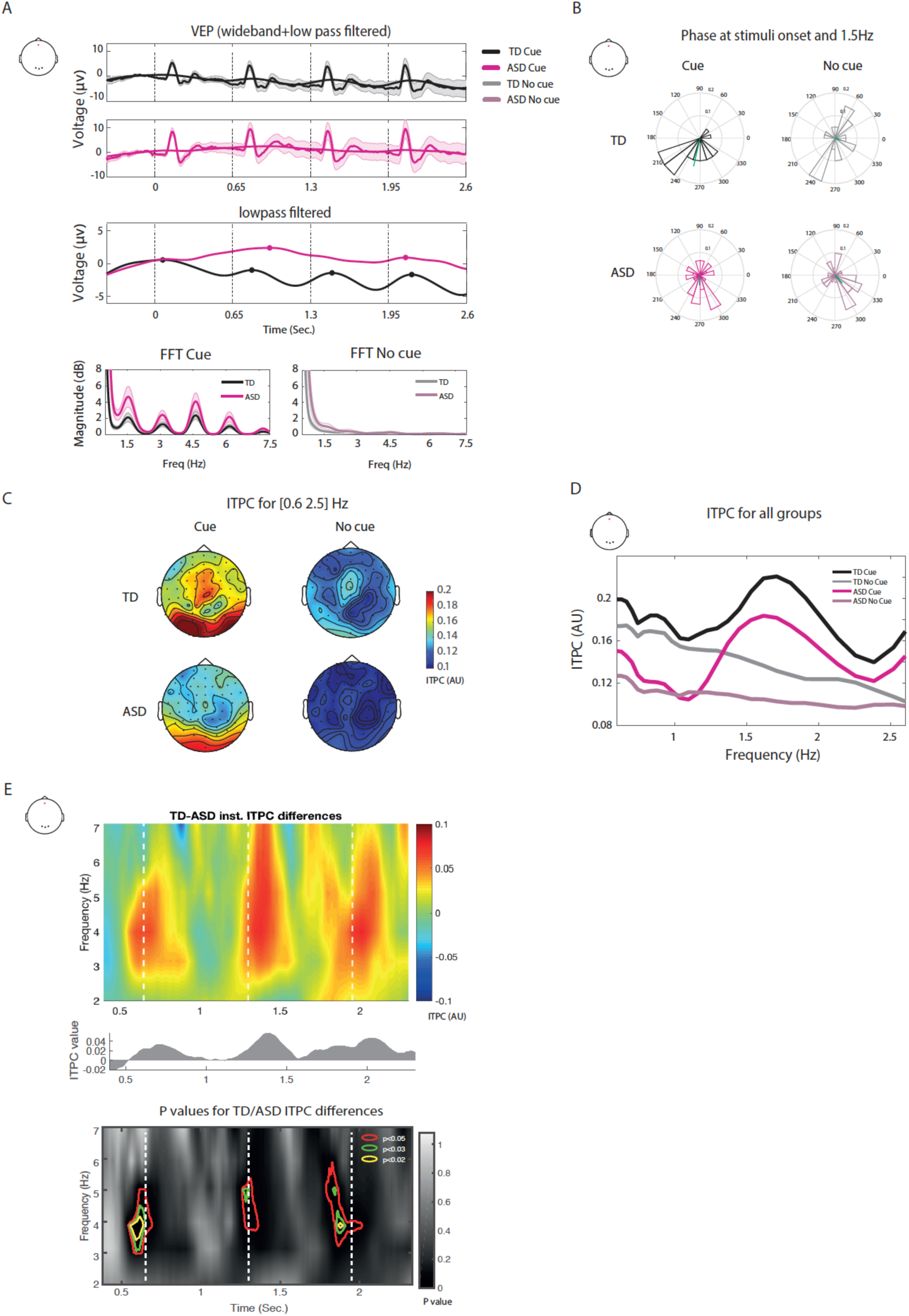
Entrainment. **A)** Top: Grand averaged low-pass filtered signals (1.9 Hz) superimposed on wideband signal, time locked to the first of the visual stimuli, for an epoch that extends through the four visual cue stimuli. Onsets of visual stimuli are marked by vertical dashed lines. Middle: Low-pass signals for ASD and TD groups. Bottom: Averaged FFT for TD and ASD groups with standard error, for the Cue (left) and No-Cue conditions (right). Note that none of the observed differences were statistically significant. **B)** Polar plots of phase concentration at 1.5 Hz at time of stimulus presentation for ASD and TD groups for Cue and No Cue conditions. **C)** Topography plots of ITPC for ASD and TD groups for Cue and No-Cue conditions. **D)** ITPC across time for 2^nd^, 3^rd^ and 4^th^ cues in the sequence. **E)** Top: ITPC for each time and frequency bin. Middle: Mean ITPC value collapsed across all frequency values at each time point. Bottom: Comparing ITPC between groups over time for different frequency bins. P-values for permutation tests (N=10000) on the group comparison. In colored contours: patches with significant values (Data interpolated for visualization). As can be seen in Figure 4E, ITPC is significantly stronger in the TD compared to the ASD group prior to the 2^nd^ and 4^th^ cue onset, and after all cue onsets in this examination (2^nd^, 3^rd^ and 4^th^ cues in the sequence). To see if behavioral measurements, and specifically a measurement of behavioral variability, correlated with neural entrainment, we measured the correlation between RT and ITPC, and standard deviation of RT and ITPC. Through this correlation, we wished to investigate whether there is a direct link between speed of processing and entrainment, and/or whether the degree of consistency of response is correlated with the consistency of neural entrainment, i.e. ITPC. While no correlation was found between ITPC and RT, an inverse linear correlation between SD-RT and ITPC at 1.5Hz was demonstrated for both the TD (Pearson ρ=-0.46, p<0.01) and the ASD groups (Pearson ρ=-0.47, p<0.01), and across the two groups (Pearson ρ=-0.46, p<0.01, see Fig. 2D). That is, in both groups, the higher the ITPC of an individual participant, the lower the variance of their RTs. To find clusters in this correlation distribution, K-means classification was performed on all participants SD-RT*ITPC correlation data. Classification using the elbow method for finding optimal number of clusters (Allen *et al.*, 2014), resulted in two clusters. Notably, 75% of the high SD-RT cluster contains data points from ASD participants; 54% of the low SD-RT cluster contains data points from TD participants. Among the ASD group, 66% of participants belong to the high SD-RT cluster, and 33% to the low SD-RT cluster. Among the TD group, 70% of participants belong to the low SD-RT cluster, and 30% to the high SD-RT cluster. Auditory Evoked Response (averaged AEP): Group*Condition Analysis of Variance (ANOVA) was performed on the auditory components P1 at 50ms, N1 at 100ms and P2 at 200ms (See Fig. 5). Analysis showed no main effects or interactions of the factors Group and Cue, over fronto-central scalp for the P1 (Group: F=1.8; df = 1; p=0.19; Cue: F=0.14; df = 1; p=0.7 ns; interaction: F=3.1; df = 1; p=0.08) or P2 (Group: F=1.95; df = 1; p=0.1 ns; Cue: F=0.4; df = 1; p=0.52; interaction: F=0.17; df=1; p=0.67). For N1, however, a main effect of Group was observed (F=44; df=1 p<0.001); a main effect of Cue was found as well (F=184; df = 1; p<0.001), although the two factors did not interact significantly (F=0.19; df=1; p=0.6).

**Figure 5:**
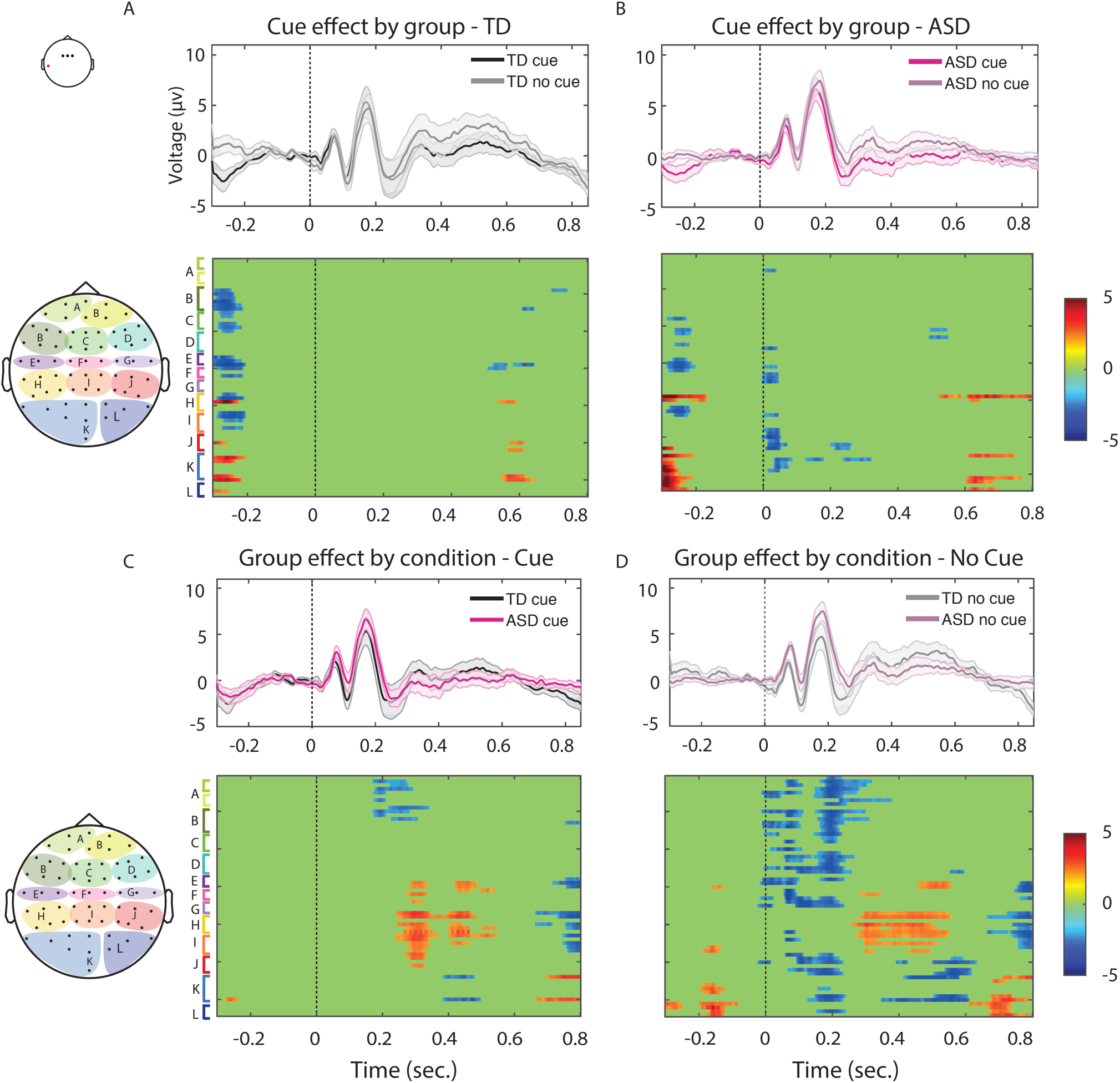
Auditory evoked potentials. Waveforms and statistical cluster plots (SCP) for Cue versus No-Cue conditions for TD and ASD groups (A and B, respectively) and for ASD versus TD groups for Cue and No-Cue (C and D, respectively) conditions. Waveforms are plotted for fronto-central channels. SCP colors show results where at least 10 consecutive time points were significant, for two adjacent channels. Note that early differences in D may be the result of baseline differences.

Finally, Pearson correlation between behavioral/cognitive assessments of the ASD children and ITPC found no significant correlations. Tests were performed for Autism Diagnostic Observation Schedule (ADOS; rho=0.06, p=0.84); Performance IQ (rho=-0.02, p=0.87); Verbal IQ (rho=0.12; p=0.83) and Repetitive Behavior Scale-Revised (RBS-R; rho=-0.17; p=0.83).

## Discussion

Clinical observations and behavioral tasks suggest that people with ASD are often inflexible in adapting to changes in their environment (Van de Cruys *et al.*, 2014), but the field lacks mechanistic insight into the neural underpinnings of this prominent feature of the Autism phenotype (Lawson *et al.*, 2017). We reasoned that one potential source might be disruption of synchronous neuronal activity at the neural circuitry level, a neural mechanism that has been implicated strongly in the formation of anticipatory predictive processes (Calderone *et al.*, 2014) and in the representation of rhythmic environmental structures (Thut *et al.*, 2012). The impairments observed in children with autism in tasks that involve adaptation to stimuli and/or anticipatory processing raise the possibility that impairment in the coherence of underlying neural activity might play a role (Coben *et al.*, 2008; Schwartz *et al.*, 2017; Foxe *et al.*, 2018).

To this end, we applied a paradigm designed to evoke neural entrainment and anticipatory processes, enabling us to study these functions in children with ASD. We presented a series of four visual isochronous cues, followed by an auditory target that occurred in rhythm with the prior cues. A control condition did not include any cues prior to the auditory target. Participants performed a speeded target detection task while high-density EEG and response speeds were measured.

Our analysis of the electrophysiological data revealed that visual evoked responses were essentially indistinguishable between the groups (Fig. 3A; Fig. 5), while anticipatory activity prior to the anticipated stimuli showed an atypical pattern in the ASD group (Fig 2). Taken together, these two observations point to intact sensory processing (Brandwein *et al.*, 2013; Foxe *et al.*, 2015; Butler *et al.*, 2017; Beker *et al.*, 2018), despite altered entrainment and anticipatory processing in this population. These results align well with a previous study by our group, where anticipatory alpha-band oscillatory activity was investigated in the context of a cued intersensory attention task (Murphy *et al.*, 2014). In that study, participants were cued on a trial-by-trial basis to prepare to respond to an upcoming target stimulus, which was presented after a delay of about 1300ms. On a given trial, the participant either prepared to respond to the auditory component of the target, or to the visual component. In this way, during the anticipatory period between cue and target, attention was deployed differentially according to the anticipated sensory task. In paradigms like this, a very consistent finding is that there is a strong increase in alpha-band activity over visual cortex when the participant is getting ready to respond to the auditory component, which has been linked to the anticipatory suppression of the visual input (see e.g. (Foxe *et al.*, 1998; Fu *et al.*, 2001)), an effect that is also readily seen in neurotypical children (Murphy *et al.*, 2016). However, in Murphy et al. (2014), this effect was significantly reduced in children with ASD, suggesting that transiently deployed anticipatory neuro-oscillatory mechanisms were weakened in this population. The current work extends these findings by showing that neuro-oscillatory activity in a lower frequency band that entrains to the rhythm of a predictable task is also impacted in ASD.

We measured anticipation by comparing amplitudes of the signals in the 200 ms period immediately prior to each of the visual cue stimuli and also just preceding the target auditory stimulus (the CNV), as well as by measuring phase coherence of the slow oscillatory activity at the frequency of stimulus presentation. In the TD group, we found the predicted gradation of CNV over frontal sites, prior to each stimulus onset. There was an evident build function to the CNV such that it increased systematically in amplitude as the cueing sequence progressed and the imminence of the target approached (Fig. 3B). This pattern was significantly weakened in the ASD group over frontal scalp. Interestingly, we observed an apparent effect of expectation in the ASD group over a focal region of occipital scalp, which was not seen in the TD group. This similarity of timing and overall function, in the face of different topographies, suggests that the two groups deployed different brain circuits to solve the same task. However, it is important to point out that this posteriorization of anticipatory processes in ASD was not predicted, has not been previously reported, and was uncovered only in post-hoc exploratory analysis. As such, it should be considered with caution and will need to be replicated. The posterior focus is consistent with a sensory-based representation of anticipatory processing in ASD, and contrasts with the typical fronto-central distribution of the CNV, which is thought to reflect motor or psychological planning prior to a pending target (Walter *et al.*, 1964; Tecce, 1972; Rosahl & Knight, 1995). Finding atypical anticipation of expected stimuli, despite intact sensory responses, aligns with one of the prominent theories in ASD - the weak central coherence theory (Happe & Frith, 2006). This theory proposes that there is a failure to extract global form and meaning in ASD, resulting in a processing bias for featural and local information, and relative difficulty in extracting gist in everyday life. Indeed, it has been argued that people with ASD might apply strategies that depend on the sensory inputs, at the expense of more integrative processing that requires an awareness of context necessary for prediction (Happe & Frith, 2006; Sinha *et al.*, 2014). The dissociation we found between visual evoked responses, which were similar between the groups, and classic anticipation-related components over anterior scalp regions, which were reduced for the ASD group, suggests intact immediate sensory evoked activity, together with altered anticipatory-related activity in the ASD group.

Further indication for reduced tracking of regularities in the ASD group comes from entrainment, as measured by phase coherence of the signal relative to a highly predictable stimulus onset. Inter-Trial Phase Coherence (ITPC) measures the consistency of phase within a selected frequency band across trials. Our data show reduced ITPC for the ASD participants at the 1.5 Hz stimulus presentation rate. Also, when calculated on each time point in the trial, a higher ITPC for the TD group was seen in the theta band immediately before the onset of the visual cues (Fig. 4E). Together, altered entrainment (ITPC) and anticipatory processing (CNV) for the ASD group point to reduced tracking of temporally predictable stimuli.

Temporal expectation often improves sensory processing (Correa & Nobre, 2008; Rohenkohl *et al.*, 2012). We observed this effect in our behavioral data, with both groups reacting faster to the Cue compared with the No-Cue condition. Entrainment of oscillatory activity may reflect a form of preparation (Lakatos *et al.*, 2008), and temporally predictable targets are associated with stronger alignment of delta phase with the target event, and with speeding of reaction time (Stefanics *et al.*, 2010). We failed to show a correlation between phase coherence and behavior. However, most studies that have identified correlation between phase in various frequency bands to behavioral performance, focused on activity preceding temporally unpredictable stimuli presented at threshold (Frund *et al.*, 2007; Fiebelkorn *et al.*, 2011; Vanrullen *et al.*, 2011) or with varying degree of expectancy (Stefanics *et al.*, 2010). Here, the mere presence of a single temporal cue may have been sufficient to improve response times to a supra-threshold target to near ceiling levels, rendering the influence of entrainment negligible. Alternatively, the relationship between phase and reaction time could be such that a monotonic regression would not reveal, for example, when a given range of phases is associated with two or more distinct RT values (Frund *et al.*, 2007; Vanrullen *et al.*, 2011). Importantly, we found that phase coherence was correlated with variability of reaction times, and this was the case for both the TD and the ASD groups. As can be seen in Figure 2C-D, along the SD-RT and ITPC dimensions, the ASD group has greater representation in the lower SD-RT/higher ITPC cluster (66% of ASD participants), while the TD group has greater representation in the higher SD-RT and lower ITPC cluster (70% of TD participants).

In accordance with their higher phase coherence and CNV in the Cue condition, we expected TD participants to show greater reaction time speeding than ASD in the cued compared to the No-Cue condition. However, both groups benefited from the predictive cues and there was Condition*Group interaction for reaction times. While not a specific focus of the current study, it is notable that the ASD group was significantly slower overall in both cued and uncued conditions. It bears emphasizing that this was not a modest difference but was found to be over 100ms, and that RT slowing in cognitive tasks is common in ASD across a range of tasks (Brandwein *et al.*, 2013; Van der Hallen *et al.*, 2015; Pirrone *et al.*, 2017; Pirrone *et al.*, 2020), and, among other changes in neuromotor functions, might represent motor control deficits in individuals with ASD (Morrison *et al.*, 2018).

Based on our data, we stress the following: in an environment where sensory events are presented in a predictable temporal pattern, neuronal responses in children with ASD are governed more by the occurrence of the events themselves, and less by their anticipated timing based on rhythm. This supports a framework whereby individuals with ASD are impaired in generating expectations based on prior experience.

A limitation of the current work is the lack of correlation between neuronal measurements and clinical measures: Although we expected to find an association between impaired entrainment and phenotypic expression of rigidity and insistance on sameness as assessed by the RBSR, this was not evident in our data. It is important to point out that the final score of an assessment reflects a weighted average of many symptoms of the disorder, while ITPC could be correlated with the severity of a specific symptom/subset, but not others. Therefore, future work that accounts for hidden patterns of relationships between ITPC and

ASD symptoms may better serve to elucidate these associations. Another potentially important consideration is our use of an inter-sensory design, whereby the auditory target is preceded by visual cues. Since there are well-documented deficits in multisensory integration in children with autism (Russo *et al.*, 2010; Brandwein *et al.*, 2013; Stevenson *et al.*, 2014a; Stevenson *et al.*, 2014c; b; Brandwein *et al.*, 2015; Foxe *et al.*, 2015; Ross *et al.*, 2015; Beker *et al.*, 2018), it is possible that impaired ITPC is due to poor cross-sensory communication and that what we are observing reflects a multisensory deficit rather than a general entrainment issue. Arguing against a pure multisensory deficit account, the ASD group showed faster RTs in the cued compared to the uncued condition, demonstrating intact use of a cross-sensory cue. An intra-sensory design (e.g., utilizing visual cues and visual targets) would address whether such entrainment and anticipation deficits are specific to a multisensory setup.

## Conclusions

High-density EEG in children with ASD indicated significantly reduced predictive processing and impaired tracking of periodic stimuli. These alterations were detected in neuro-oscillatory frequencies tracking the rhythm of the task stimulation. This implies that entrainment of neural oscillations, necessary for cognitive processes that are built upon precise timing of neuronal response like tracking and predictive processing, may be compromised in ASD.

## Author Contributions

SM and JJF conceptualized the study. SB coordinated and oversaw study implementation, and participated in the data collection. SB, JJF and SM consulted closely with each other during the data analysis and statistical testing of said data. SB carried out these analyses and made the illustrations. SB then wrote the first substantial draft of the manuscript. The three authors all contributed to the subsequent editing of the manuscript. All authors approve this final version for publication.

## Funding

Work by the authors on autism spectrum disorder is primarily supported through an RO1 from the NICHD (HD082814). Our work on developmental disabilities is also supported in part through the Rose F. Kennedy Intellectual and Developmental Disabilities Research Center (RFK-IDDRC), which is funded through a center grant from the Eunice Kennedy Shriver National Institute of Child Health & Human Development (NICHD U54 HD090260; previously P30 HD071593). Drs. Foxe and Molholm’s work in ASD also receives support through pilot grant funds from The Harry T. Mangurian Jr. Foundation.

## Data Sharing

The authors will make the full de-identified dataset with appropriate notation and any related analysis code available in a public repository (Figshare) and include digital object identifiers within the final text of the paper, so that any interested party can access them.

## Abbreviations

ASD: (Autism Spectrum Disorder)
TD: (Typically Developed)
EEG: (Electroencephalogram)
CNV: (Contingent Negative Variation)
ITPC: (Inter-Trial Phase Coherence)
RT: (Reaction Time)
SD: (Standard Deviation)
df: (degrees of freedom)
ANOVA: (Analysis of Variance)

## Conflict of interest statement

All authors confirm no competing financial interests.

## Notes

### Competing Interest Statement

The authors have declared no competing interest.

